# Analysis of amino acid change dynamics reveals SARS-CoV-2 variant emergence

**DOI:** 10.1101/2021.07.12.452076

**Authors:** Anna Bernasconi, Lorenzo Mari, Renato Casagrandi, Stefano Ceri

## Abstract

Since its emergence in late 2019, the diffusion of SARS-CoV-2 is associated with the evolution of its viral genome^1,2^. The co-occurrence of specific amino acid changes, collectively named ‘virus variant’, requires scrutiny (as variants may hugely impact the agent’s transmission, pathogenesis, or antigenicity); variant evolution is studied using phylogenetics^3–6^. Yet, never has this problem been tackled by digging into data with ad hoc analysis techniques. Here we show that the emergence of variants can in fact be traced through data-driven methods, further capitalizing on the value of large collections of SARS-CoV-2 sequences. For all countries with sufficient data, we compute weekly counts of amino acid changes, unveil time-varying clusters of changes with similar – rapidly growing – dynamics, and then follow their evolution. Our method succeeds in timely associating clusters to variants of interest/concern, provided their change composition is well characterized. This allows us to detect variants’ emergence, rise, peak, and eventual decline under competitive pressure of another variant. Our early warning system, exclusively relying on deposited sequences, shows the power of big data in this context, and concurs to calling for the wide spreading of public SARS-CoV-2 genome sequencing for improved surveillance and control of the COVID-19 pandemic.

## Main

The fast emergence of SARS-CoV-2 mutations and their possibly severe epidemiological implications call for continuous and worldwide monitoring of viral genomes. Several organizations offer repositories for depositing sequences; the largest collection is provided by GISAID^7^, which recently hit two million deposited records. Genomic surveillance concentrated initially on the monitoring of individual amino acid changes, such as the rise to dominance of the D614G Spike mutation worldwide^1^ or the spread of the A222V Spike mutation from Spain throughout Europe during the 2020 Summer^6^. As the COVID-19 pandemic progressed, research interests shifted towards the study of coordinated mutations. The term ‘variant’ has come into common use, even among the general public, to denote a set of such co-occurring amino acid changes^8^. Emerging variants have been playing an important role in the course of the pandemic, as they have been associated with increased transmission rates and changed antigenicity of the SARS-CoV-2 virus, possibly hampering testing, treatment, and vaccine development^9–13^.

The definition of variants is produced by phylogenetic analysis^3–6^. Phylogenetic trees describe the precise chain of evolutionary changes that leads from one sequence to the next, resulting in a powerful separation of viral sequences into clades or lineages that share common ancestries—henceforth, the same amino acid changes. A number of conventions and nomenclatures are in place to name SARS-CoV-2 strains based on phylogenetic guidelines (e.g. Pangolin^14^, Nextstrain^15^, GISAID^7^). The World Health Organization (WHO) has recently proposed a rapidly and widely accepted naming scheme for variants, based on the Greek alphabet^16,17^, which – as of June 3rd, 2021 – recognizes four variants of concern (Alpha^18^, Beta^19^, Gamma^4^, and Delta^20^) and seven variants of interest (Epsilon^21^, Zeta^22^, Eta^23^, Theta^24^, Iota^25^, Kappa^26^, and Lambda^27^). Several national and international organizations offer surveillance of variants and their effects^17,23,28,29^.

### A big data approach to scout variants

Here, we propose a data-driven method exclusively relying on the analysis of sequences collected in repositories. The method aims to trace the emergence of variants in specific regions (e.g. countries) using time-series of amino acid change prevalence, defined as the fraction of genome sequences with a specific change. For each country, from January 2020 until the beginning of June 2021, we counted how many sequences present given amino acid changes weekly. United Kingdom has the biggest dataset, with 3,808 changes collected over 64 weeks, whereas South Africa has the smallest (84 changes, 56 weeks). We observed that, on emergence, the prevalence of changes that will eventually characterize a variant typically show an exponential-like temporal growth^30^; one meaningful example is shown in Extended Data Fig. 1.

We harnessed the peculiarity of these temporal trends to scout emerging variants. To that end, we applied standard time-series clustering techniques to group together changes showing similar prevalence behavior over a one-month-long period. We regarded as warnings of possible variant emergence those clusters characterized by (i) a positive trend in the prevalence time-series of the constituting amino acid changes, and being sufficiently different from clusters that caused previous warnings (to avoid extracting twice a set describing the same candidate variant). To assess whether these clusters successfully match known variants (named ‘hit’), we tested their similarity against the characterizing changes of the most widespread SARS-CoV-2 lineages, as defined by Pangolin, collected in the ‘lineage dictionary’ (see Extended Data Tables 1 and 2). We also used community analysis to assess whether and how the emergence of new variants can lead to a reorganization in the dynamics of amino acid changes, as observed through their co-occurrence within clusters.

Our approach must not be considered an alternative to phylogenetic analysis, which takes into account the entire evolutionary history of the viral genome. Other methods have previously complemented phylogenesis, namely by describing typical SARS-CoV-2 mutational profiles across different countries and regions^31,32^, proposing statistical indicators for location-based mutation evolution^33^, and observing changes that become recurrently prevalent in different locations, thus suggesting selective advantages^1^. Time-based analyses have been considered for phenetic clustering of prevalent SARS-CoV-2 mutations over time^34,35^, trend detection in SARS-CoV-2 short nucleotide sequences^36^, and single amino acid changes^37^.

### Searching variants by mining time-series

To show our methods in action, Fig. 1 displays the case of Japan, featuring a matrix of 594 amino acid changes observed over 63 weeks. We identify 132 clusters, ten of which are included in the early warning system, tracking the emergence of possible variants. Four of these clusters revealed to be early hits, i.e. showing strong cluster-lineage similarity at the time of clustering. These hits occur on Jun-wk4-20, Aug-wk2-20, and March-wk3-21 (two instances), respectively associated with variants JP2, JP1, and JP3/Alpha. We use JP1, JP2, and JP3 for lineages B.1.1.214, B.1.1.284, and R.1 since, according to Pango lineages reports^38^, they originated in Japan. Each hit is qualified by the growth of the change prevalences in the identified cluster (left panels a–c) and by the similarity between cluster composition and the lineage dictionaries (central panels a–c). Partial overlaps of identified clusters with more than one dictionary are due to overlaps between dictionaries. We find that the emergence of a new variant is typically associated with a drastic reorganization of the within-cluster co-occurrence patterns of amino acid changes (right panels a–c), which may suggest profound effects of variant emergence. Panels d–g of Fig. 1 show the temporal dynamics of the changes associated with each identified variant. Strong cluster-dictionary matches, illustrated by circle size and color, are found for all variants, spanning several months after the moments when warnings are issued. By contrasting variant dynamics with recorded cases (shaded bars), we observe that JP2, JP1, and Alpha prevalences peaked ahead of the first, second, and third COVID-19 waves in Japan. By contrast, JP3 shows a prevalence peak when reported cases were at a minimum. Taken all together, Fig. 1 constitutes a syntactic fingerprint of how variants have evolved in Japan. In addition to these four hits, our method identified other six candidates that could be rapidly dismissed, because clusters identified in subsequent weeks showed low similarity with the original candidate. Four of them were discarded two weeks after the warning, the other two within five weeks. Extended Data Fig. 2 shows all ten warnings.

**Figure 1.**
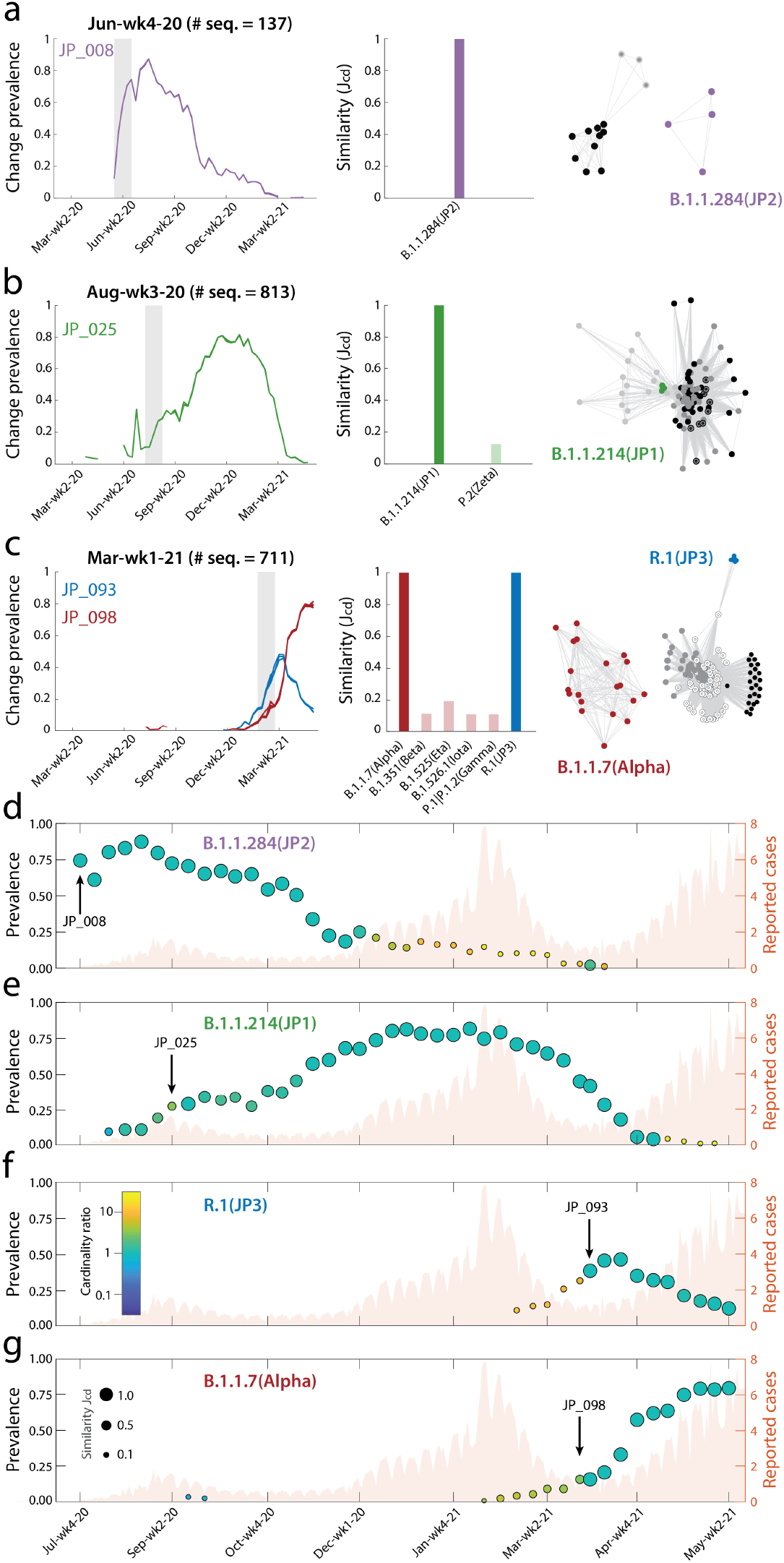
Data-driven identification of variants in Japan. **a–c**, Left panels: Four hits in Japan (two of which occurring in the same week), corresponding to clusters JP_008 (Jun-wk4-20), JP_025 (Aug-wk3-20), JP_093, and JP_098 (both observed in Mar-wk1-21). Central panels: Cluster-dictionary similarity (*J_cd_*) for the four early hits showing, at time of detection, to strongly match (*J_cd_* = 1 in all cases) the four lineages B.1.1.284 (JP2), B.1.1.214 (JP1), R.1 (JP3), and B.1.1.7 (Alpha); values *J_cd_* < 0.1 are not shown. Right panels: Community detection applied to the within-cluster change co-occurrence matrix evaluated over the period of time between the emergence of two subsequent variants. The resulting interaction network is plotted using a force-directed layout^41^, with edge lengths inversely proportional to link weights. Colors indicate nodes belonging to communities that strongly match known lineages, other communities are in gray-scale shades. **d–g**, Temporal dynamics of the four identified variants. In each scatter plot, the left y-axis value represents the average prevalence of the changes in the cluster-dictionary intersection, circle size represents the cluster-dictionary similarity (*J_cd_* < 0.1 not shown), while circle color is proportional to the log-ratio between the number of changes in the cluster and in the dictionary. Warnings are marked with a labeled arrow. In the background, the number of reported COVID-19 infections (thousand cases, right axis) in Japan^42^.

### Variant emergence around the world

Fig. 2 illustrates how our method reconstructs the emergence of the main WHO-named variants at their country of origin. The temporal pattern of the Alpha variant is remarkably neat, exhibiting an initial phase of exponential growth, and well correlating with the number of reported COVID-19 cases. After a ten-weeks plateau with prevalence values close to 100%, a five-weeks sharp decline occurred, associated with the concurrent growth of the Delta variant (see Extended Data Fig. 3). Note that some variants never reached a high prevalence, e.g. Epsilon and Iota. Variant dynamics also differed in terms of peak timing. For instance, variant Zeta well matched the temporal evolution of the second COVID-19 wave in the country, and Gamma shortly anticipated the third large wave of infections and became locally dominant. Similarly, variants Kappa, Delta, and Beta, grew quite significantly before the second wave of cases in their respective countries of origin. Note that some weaker matches are found also before the relevant warnings, essentially because some amino acid changes included in the lineage dictionaries did indeed emerge individually well before the diffusion of variants. Also, note that the WHO-named variants Eta, Theta, and Lambda do not appear in this figure as their countries of origin did not meet our criteria for minimum data availability and were not included in the analysis.

**Figure 2.**
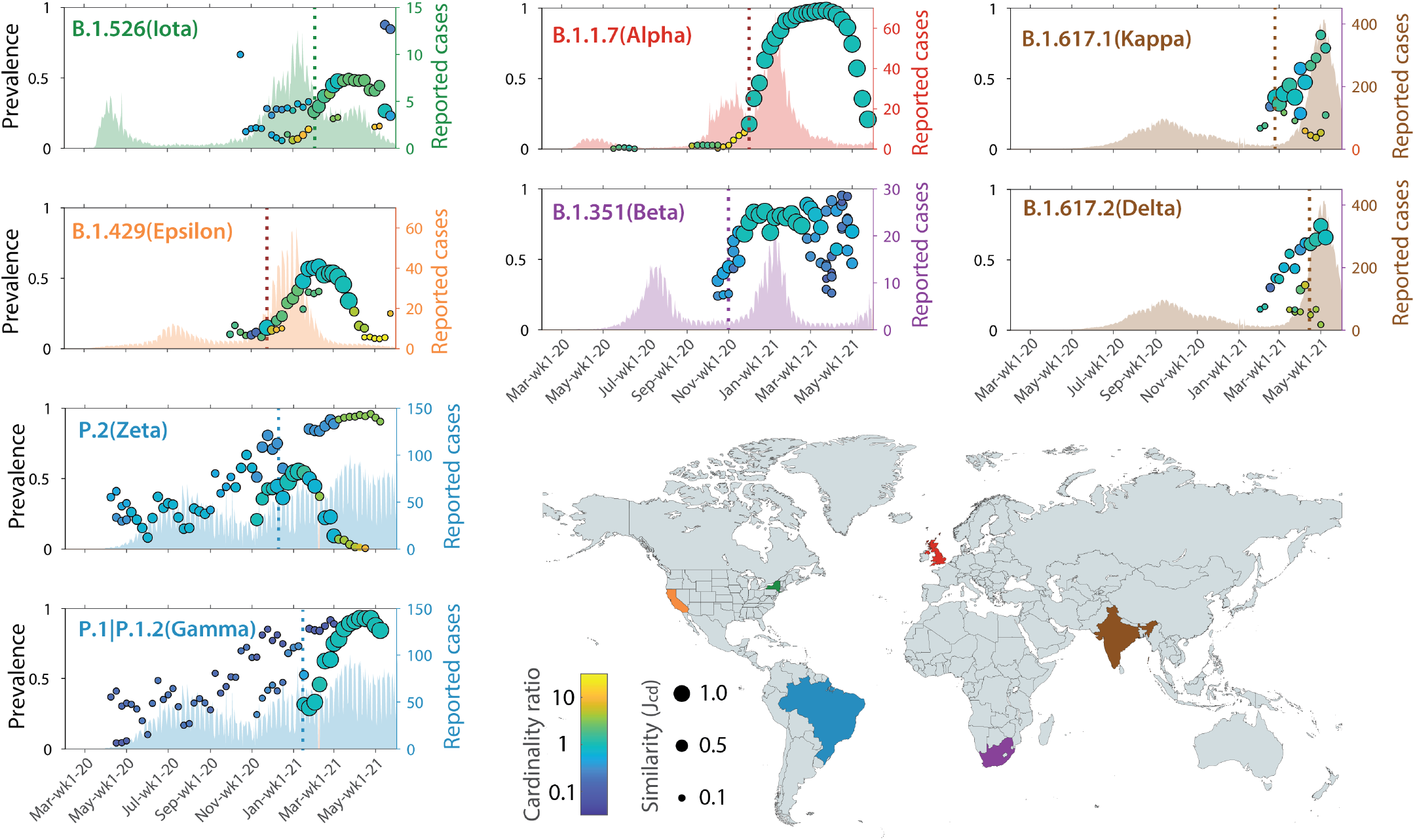
Emergence of variants in their country of origin. Details of the variant dynamics plots shown within insets as in Fig. 1d–g. The vertical dashed lines indicate the dates of hits using our method. Estimates of reported cases are taken from Johns Hopkins/Our World in Data^42^, except for the US, for which data comes from the Centers for Disease Control and Prevention^43^.

Fig. 3 shows the temporal patterns of notable variants as identified by our method in Europe and in the US. For all European countries with at least 16k deposited sequences, panel a displays the growth curves of changes associated with the Alpha variant at the time when the relevant warnings were issued by our method. The spread of Alpha outside the UK was delayed by five to seven weeks; warnings concentrate on three consecutive weeks in late January (country shading). Interestingly, the growth curves of the change prevalences associated with the Alpha variant outside the UK are more noisy than inside the UK, likely due to availability of fewer sequences and spatial heterogeneities in sequence deposition.

**Figure 3.**
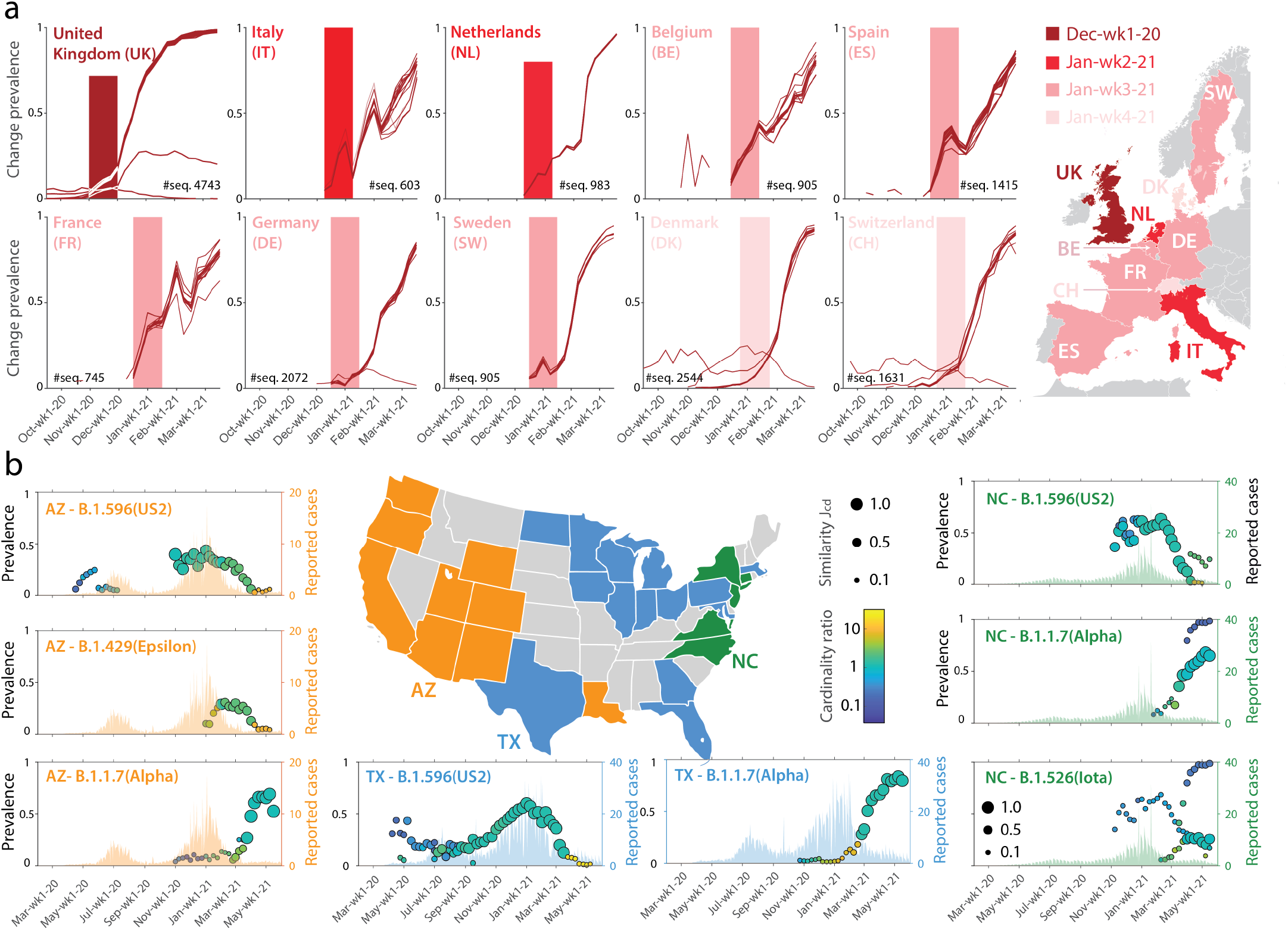
Temporal dynamics of notable variants in European countries (panel a) and in the US (b). Color shading for European countries indicates the timing of emergence of the Alpha variant (the lighter, the later). Different colors code instead the couple/triple of variants (Epsilon, Iota, Alpha, and US1/US2 as aliases of B.1.2 and B.1.596— only US2 is shown here) that were detected and tracked in the US using our method. Inset details as in Fig. 1d–g.

Panel b of Fig. 3 shows instead the dynamics of the most widespread US-born SARS-CoV-2 variants in all US states where at least 5k sequences were deposited and their interplay with WHO-named variants. Our clustering and warning methods partitioned these states into three distinct, geographically quite compact, groups. A central/eastern group of states (blue) displays a baseline temporal pattern where common US variants first rose until Jan-wk1-2021, then started declining under pressure of variant Alpha. In the Western group of states (orange), the Californian-born variant Epsilon appeared in-between the US2 variant and Alpha, whereas in the north-eastern group (green) the New-York-born Iota appeared after the rise of Alpha, none of them reaching the same prevalence as Alpha. Although quite general, the patterns above described in groups may show country-dependent peculiarities. In Louisiana, for example, the hit for Epsilon followed the one for Alpha, and we did not issue warnings for Alpha in New York, nor for US1/US2 in New Jersey and Connecticut.

## Discussion

We have presented a novel methodology to scout SARS-CoV-2 variants entirely based on the analysis of data from the GISAID sequence repository; specifically, we have shown that emerging lineages can be detected via cluster analyses of the prevalence time-series of amino acid changes. Our scouting method is not in competition with phylogenetic analysis, which is fundamental for properly defining variants as lineages through monitoring of the evolution of viral sequences. Nevertheless, the proposed big-data-driven approach proved to be quite effective. First, the major WHO-named variants of SARS-CoV-2 were well and timely captured in their country of origin, an indication that the early dynamics of change prevalence is a distinguishing property of emerging variants. All hits were fast (requiring no more than five weeks, the length of the time window used for cluster analysis), except the hits of Beta (five plus two weeks) and Iota (five plus one weeks)—see Table 1. Second, our plots reveal that the method can effectively trace and visualize the variants’ natural evolution. Variants emerge, reach their maximum prevalence, and eventually start a declining phase, typically when a competing variant emerges. Our method also issues warnings that are not found to match major lineages down the road. We do not see this outcome as an intrinsic limitation of our approach, though. In fact, even when not selected as hits, warnings are informative of the dynamics underlying variant emergence and inter-variant competition; in addition, similarity analysis can clarify, typically in a matter of a few weeks at most, whether an emerging variant is gaining traction.

**Table 1.**
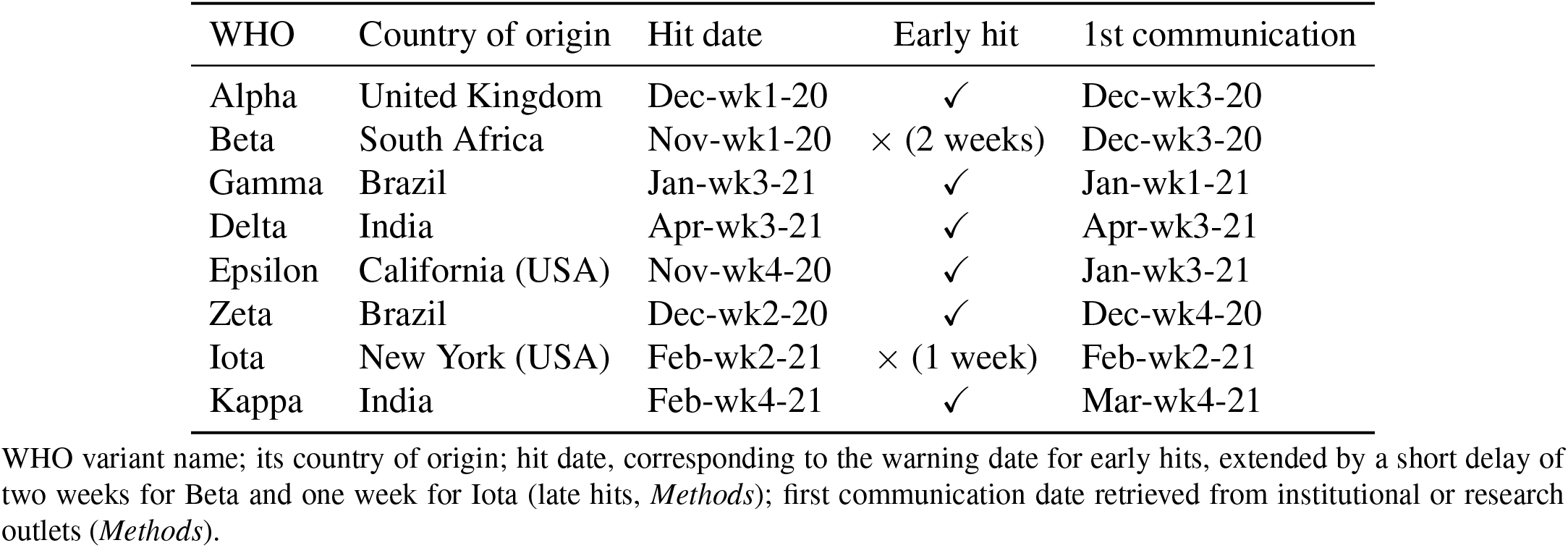
Capture of the emergence of the major WHO-named SARS-CoV-2 variants

Relying exclusively on big data for variant scouting is hampered by three problems: (i) consistency of sampling, (ii) delay of deposition, and (iii) biases in sampling. As for (i), the ratio between the number of sequences and reported COVID-19 cases widely varies among countries and drops from 77% in Iceland (clearly facilitated by small number of cases) to below 0.1% for many countries, also including some large ones like India and Brazil. Among the countries with lots of cases, the UK stands with an exceptionally high ratio exceeding 9%. Indeed, US and UK have contributed the largest number of sequences worldwide (517k and 425k, respectively). Extended Data Table 3 shows these statistics for all countries contributing to GISAID with more than 1,000 sequences, whereas Extended Data Table 4 shows statistics for the 50 US states. Regarding (ii), as an example, in the UK the average delay between collection and deposition amounts to 24 days. This delay tended to reduce as the pandemic unfolded, from 38 days in 2020 to just 16 days in 2021. Iceland is again striking the best performance, with 11 days of average delay in 2021. Concerning (iii), heterogeneities in surveillance may introduce biases that should be considered, for instance when sampling is concentrated within regions where variants have newly appeared, thereby causing possible over-estimation of the observed prevalence of variants within a country. Another aspect interfering with a correct estimation of variants’ prevalence is the concurrent rollout of COVID-19 vaccines, particularly in case of variants possibly endowed with partial immune escape potential, as in the recent case of Delta variant^39^.

Being aware of these obstacles, the potential of developing and using big-data-driven approaches to keep track of variant emergence may be proved by comparing the timings of warnings issued by our method for the eight major WHO-named variants (as of June 3rd, 2021) with the dates of their initial communication in the media or in relevant scientific outlets (see again Table 1 and *Methods*). In five cases (Alpha, Beta, Epsilon, Zeta, Kappa), our data-driven warnings anticipated the press notifications, in some cases (Beta, Epsilon, Kappa) with a lead time of several weeks. In other two cases (Delta, Iota), the warning would have been issued around the same time as the first communications. Only in the case of the Gamma variant our warning was late (by a couple of weeks) compared to its first communication. It must be remarked that Brazil, where both Gamma and Zeta variants originated, was sequencing a low number of samples at that time.

In conclusion, our warnings were issued at times which favourably compare with most press/research announcements, in some cases even anticipating them. The above figures illustrating variant hits and dynamics collectively provide an effective fingerprint of the location-specific temporal evolution of variants, and well support comparative displays in time and space. Our exercise thus strongly corroborates the urgent call, raising from the entire scientific community, for faster and wider public publishing of viral sequences^40^.

## Methods

### Data collection from the GISAID data source

Since January 5, 2020, when the complete genome sequence of SARS-CoV-2 was first released on GenBank (Access number: NC_045512.2), there has been a rapid accumulation of viral sequences. Many tools are available for quickly extracting the assignment of sequences to lineages or the prevalence of each amino acid change in specific geographical areas, including ViruSurf^44^, `outbreak.info`^45^, CoVariants^46^, and `cov-lineages.org` lineage report^38^. Here, we considered the protein-level mutations (hereon named ‘amino acid changes’) of 1,819,996 complete SARS-CoV-2 genome sequences from infected individuals from all over the world that were downloaded on June 3rd, 2021, using EpiCov^*TM*^ data from the GISAID database (https://www.gisaid.org/^7^) on the basis of a specific Data Connectivity Agreement. The original data file contains: sequence accession ID, collection date, submission date, Pangolin lineage, colletion location, and the list of amino acid changes. All records not reporting the day (but only year, or year and month) in the collection date field were discarded, resulting in a dataset of 1,763,923 records (about 97% of the initial size). To denote amino acid changes, we adopt the common notation used by GISAID, representing them as the concatenation of the following elements: the protein acronym, the reference amino acid residue, the position (using the coordinate system of the single protein), and the alternative amino acid residue exhibited by the specific mutated sequence (in case of substitutions) or a dash (in case of deletions). Proteins include, for example, Spike, Envelope (E), Membrane (M), Nucleocapside (N), as well as non-structural proteins forming the ORF1ab polyprotein (NSP1, NSP2, …, NSP16). Changes can occur at any point within a protein; they are evaluated with respect to the reference sequence WIV04 (used by GISAID^47^).

### Temporal aggregation of sequence mutations

Original collection dates were temporally binned into four approximately weekly periods per month, namely days 1–7 (shortcut as wk1), 8–15 (wk2), 16–23 (wk3), and 24–end of month (wk4). We verified that the small differences in bin sizes among the four periods do not have any impact on the results obtained using our data mining methods, as they rely on change prevalence (relative count of sequences) rather than change abundance (absolute count). We transformed the GISAID data into a table containing tuples of the following kind: amino acid change, country of collection, week of collection, absolute count of sequences holding that change, and total collected sequences in the same country-week. As locations of interest, we selected all European countries and US states for which at least 16k and 5k sequences were available, respectively, and a small number of other countries where variants of concern/interest have originated according to the World Health Organization^17^. For each of those countries, we prepared a matrix where: each row *r* represents an amino acid change that has been observed in at least five sequences and for at least three weeks, each column *c* represents an observation week, each cell ⟨*r, c*⟩ contains the number of sequences exhibiting the amino acid change *r* observed in week *c*. The full account of matrix sizes for the selected countries is shown in Extended Data Table 5. We also extracted from the GISAID dataset the total number of sequences collected at given locations for each of the considered weekly periods. Data extraction and aggregation was performed using PostgreSQL (Version 12.5) and the Pandas library (Version 1.2.1) of Python (Version 3.8.5). The steps of data extraction and aggregation are summarized in Extended Data Fig. 4.

### Construction of lineage dictionary

We extracted all lineages that are expressed in at least 5k sequences worldwide and a small number of lineages that deserved attention by the news and/or other online sources, provided they appeared in at least 1k sequences worldwide. The assignment of amino acid changes to lineages is not uniquely/uniformly defined by all sources; following the choice made by Mullen *et al.*^45^, we computed the set of characterizing changes, pragmatically defined as those that appear in at least 75% of the lineage sequences in GISAID, for all retained lineages. This rule produces partially overlapping lists of amino acid changes, with lengths ranging from five to 24, which we regard as our ‘lineage dictionary’ (Extended Data Table 2). To improve readability, the naming convention starts with the Pangolin lineage^14^, followed by the WHO name^17^ (using Greek alphabet) when available. In selected cases, we add a geographical characterization, based on the country of origin (https://cov-lineages.org/^38^), resulting into US1/US2 and JP1/JP2/JP3 aliases. Only exceptionally we merged multiple lineages into one label, namely when lineages share the same set of characterizing amino acid changes. Extended Data Table 1 contains a comprehensive list of lineages considered in our study.

### Cleaning of too frequent, confounding changes

We found that few changes (globally six, four worldwide – S_D614G, NSP12_P323L, N_R203K, and N_G204R – and two specific to North-America—NS3_Q57H and NSP2_T85I, out of a total of 78,650 recorded changes) are disproportionately frequent in the dataset of each continent, being present in more than 60% of GISAID sequences and populating the dictionary of more than 30% of the most frequent lineages (i.e. those represented by at least 80 sequences). To avoid confounding effects in data analysis, we removed these changes from both the aggregated data matrices and the dictionary entries.

### Cluster analysis

To assess whether the temporal patterns of changes’ prevalence occur in a coherent manner in the dataset, as expected in the case of changes belonging to closely related variants, we apply a relatively simple time-series clustering algorithm. All the following analyses have been performed using MATLAB R2021a. At each time *t*, we retain the *n*(*t*) time-series of change prevalence (current ratio between change counts and total counts of changes) that are observed continuously over a time interval of four weeks prior to *t* ([*t* − *w* + 1 · · · *t*], with *w* = 5) and partition them via *k*-medoids clustering^48^ (PAM algorithm, `kmedoids` function in MATLAB), with pairwise distances between time-series being evaluated via dynamic time warping^49^ (`dtw` in MATLAB). The optimal value of *k* is exhaustively searched over the range [1 … min(20*, n*(*t*)*/*4)] and is selected as the one that maximizes the average silhouette score^50^ evaluated over the entire set of *n*(*t*) change prevalence time-series.

### Our early-warning system for variant emergence

To identify clusters of changes characterized by an increasing trend in their prevalence time-series, as expected in the case of emerging variants, we use the non-parametric Kendall’s *τ_B_* statistic^51^, as implemented in the `ktaub` MATLAB package^52^, namely to evaluate the ordinal association between change prevalence and sampling time. More precisely, a cluster of changes is considered to deserve further attention as a candidate for variant emergence if its average prevalence time-series shows a positive trend (*τ_B_* > 0 at significance level *α* = 0.05). Among these candidate clusters, we are particularly interested in those that are sufficiently different from previously observed trending clusters, because they could signal the emergence of a new virus variant. Indeed, the partitioning of change time-series is performed independently at each time step, thus the clusters identified at a given time are in principle not related to those identified at any previous step. However, because of the temporal autocorrelation of change prevalence dynamics, similarities are expected (and found) to exist in the composition of clusters identified at subsequent time steps. To quantify similarity for each pair of clusters, we use the classical Jaccard index^53^, defined as the ratio *J_cc_* between the cardinality of the intersection and the cardinality of the union of the sets of changes constituting the two clusters. Among the candidate clusters with a positive trend in their prevalence time-series, only those that are sufficiently different from any previous candidates (*J_cc_* < 0.5) are selected to form an early warning system for monitoring the possible emergence of new variants.

### Cluster-dictionary comparison

We use the similarity between the composition of early warning clusters and the lineage dictionaries to assign a posteriori the observed change time-series to known lineages, thereby building a ground truth for assessing the performance of data analysis. Specifically, we use again Jaccard similarity index, this time applied to cluster vs. dictionary change composition (*J_cd_*). A threshold *J_cd_* > 0.5 is used to identify clusters for which there is a close compositional match to a known lineage. If a match exists, we conclude that the method has identified the lineage (thus, the underlying variant) at the same time as when clustering occurred (early hit). If not, we keep monitoring the cluster-dictionary similarity *J_cd_* in the following time steps, each time using the cluster with the highest compositional similarity *J_cc_* with that of the previous step, provided that the inter-step similarity remains sufficiently strong (*J_cc_* > 0.5), until the threshold *J_cd_* = 0.5 is possibly exceeded (late hit). If the inter-step similarity test fails without producing a late hit, the candidate cluster causing the early warning is discarded. Rapid convergence towards a decision between late hit or candidate discarding constitutes a desirable property of our method.

The study of the changes associated with the successfully identified variants can be deepened by analyzing their temporal dynamics along with the country-specific epidemiological patterns of the pandemics. Specifically, we match the dictionary composition of each identified variant to cluster composition over time and evaluate: i) the average prevalence at the time of clustering of the changes in the intersection between cluster composition and lineage dictionary; ii) the cluster-dictionary similarity *J_cd_*; and iii) the log-ratio between cluster and dictionary cardinalities. This methodology is applied to the whole dataset, i.e. also prior to the emergence of variants or after their possible disappearance; in fact, some changes belonging to a lineage dictionary may be individually present in the population before/after the diffusion of the relevant variant.

### Assessment of the early warning system

We collected information about the first Institutional Communications (ICs), Research Communications (RCs) captured at their first version, and Published Papers (PPs) about the most important WHO-named variants to assess whether the points in time when warnings are issued by our method compare well with the points in time when variants actually became known. The following references provide the dates indicated in Table 1:

- Alpha: IC on 18-Dec-20^23^ (Technical Briefing 1 of Public Health England, PHE); RC on 19-Dec-20^54^ (post on Virological.org forum); PP by Volz *et al.*^18^.
- Beta: IC on 18-Dec-20^55^ (Republic of South Africa Health Department); RC on 22-Dec-20^56^; PP by Tegally *et al.*^19^.
- Gamma: IC on 06-Jan-21^57^ (National Institute of Infectious Diseases of Japan); RCs on 11-Jan-21^58^ and 12-Jan-21^59^ (two independent posts on Virological.org forum); PP by Naveca *et al.*^4^.
- Delta: IC on 21-Apr-21^23^ (Technical Briefing 10 of PHE); RC on 28-Jun-21^20^.
- Epsilon: IC on 17-Jan-21^60^ (California Department of Public Health); RC on 20-Jan-21^61^; PP by Zhang *et al.*^21^.
- Zeta: IC on 13-Jan-21^23^ (Technical Briefing 8 of PHE); RC on 26-Dec-20^62^; PP by Voloch *et al.*^22^.
- Iota: IC on 10-Mar-21^23^ (Technical Briefing 7 of PHE); RCs on 15-Feb-21^25^ by Caltech group and 25-Feb-21^63^ by Columbia University group.
- Kappa: IC on 24-Mar-21^64^ (Indian Ministry of Health and Family Welfare); RC on 24-Apr-21^26^.

### Community analysis

The points in time when lineage dictionaries are for the first time successfully (*J_cd_* > 0.5) matched to cluster composition are used to partition temporally the dataset of change prevalence. For each resulting time window, we explore the within-cluster co-occurrence dynamics of the different changes by building a weighted, undirected graph where the nodes are the different changes and the edges represent pairwise interactions between changes; edge labels are evaluated as the number of clusters in which two changes occur together^65^. To assess whether changes can be grouped into densely connected sets, thereby demonstrating robust co-occurrence patterns within the clusters identified over the time window of interest, we apply the Louvain method of community detection^66^ using the MATLAB function `GCModulMax1` (Community Detection Toolbox^67^). The change compositions of the resulting communities are compared with the lineage dictionaries by using again Jaccard index.

## Data availability

All data matrices used in this study, one for each country or US state, as well as the lineage dictionary, have been deposited in Zenodo at https://doi.org/10.5281/zenodo.5090169. Original sequence metadata and amino acid changes are publicly accessible through the GISAID platform.

## Code availability

The Zenodo repository https://doi.org/10.5281/zenodo.5090169 includes custom scripts that can be used to reproduce the results presented in this study.

## Acknowledgements

We gratefully acknowledge all data contributors, i.e. the Authors and their Originating Laboratories responsible for obtaining the specimens, and their Submitting Laboratories that generated the genetic sequence and metadata and shared via the GISAID Initiative the data on which this research is based. A.B. and S.C. acknowledge the support provided by the ERC Advanced Grant 693174 “Data-Driven Genomic Computing (GeCo)”.

## Author contributions

A.B. and S.C. proposed the study, all authors jointly conceptualized and designed it; A.B. and S.C. designed the data extraction methods (implemented by A.B.); R.C. and L.M. designed the data clustering and analysis methods (implemented by L.M.); A.B. mined and curated published SARS-CoV-2 variant data; all authors wrote and revised the manuscript.

## Additional information

**Correspondence and requests for materials** should be addressed to A.B.

---

**Extended Data Figure 1.**
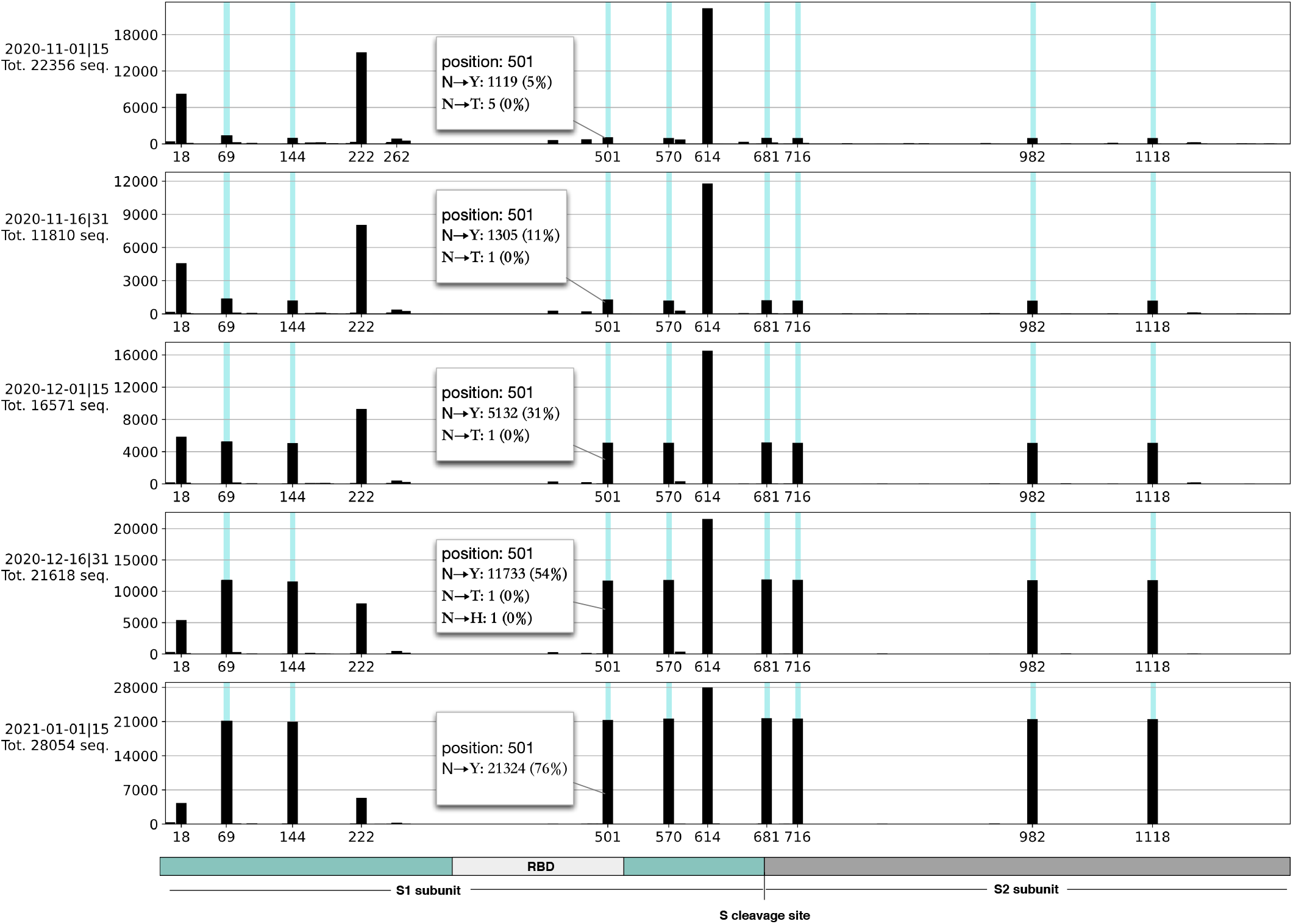
Counts of protein Spike amino acid changes in the UK in different bi-weekly periods. The tracks represent sequences of the five groups filtered by date (from the first half of November 2020 till the first half of January 2021). A blue background is used in the positions notably belonging to the Alpha variant, as first recognized as the B.1.1.7 lineage by Rambaut *et al.*^54^: H69-, V70-, Y144-, N501Y, A570D, P681H, T716I, S982A, and D1118H; higher black bars in these positions from top to bottom indicate an increase in the frequency of these amino acid changes.

**Extended Data Figure 2.**
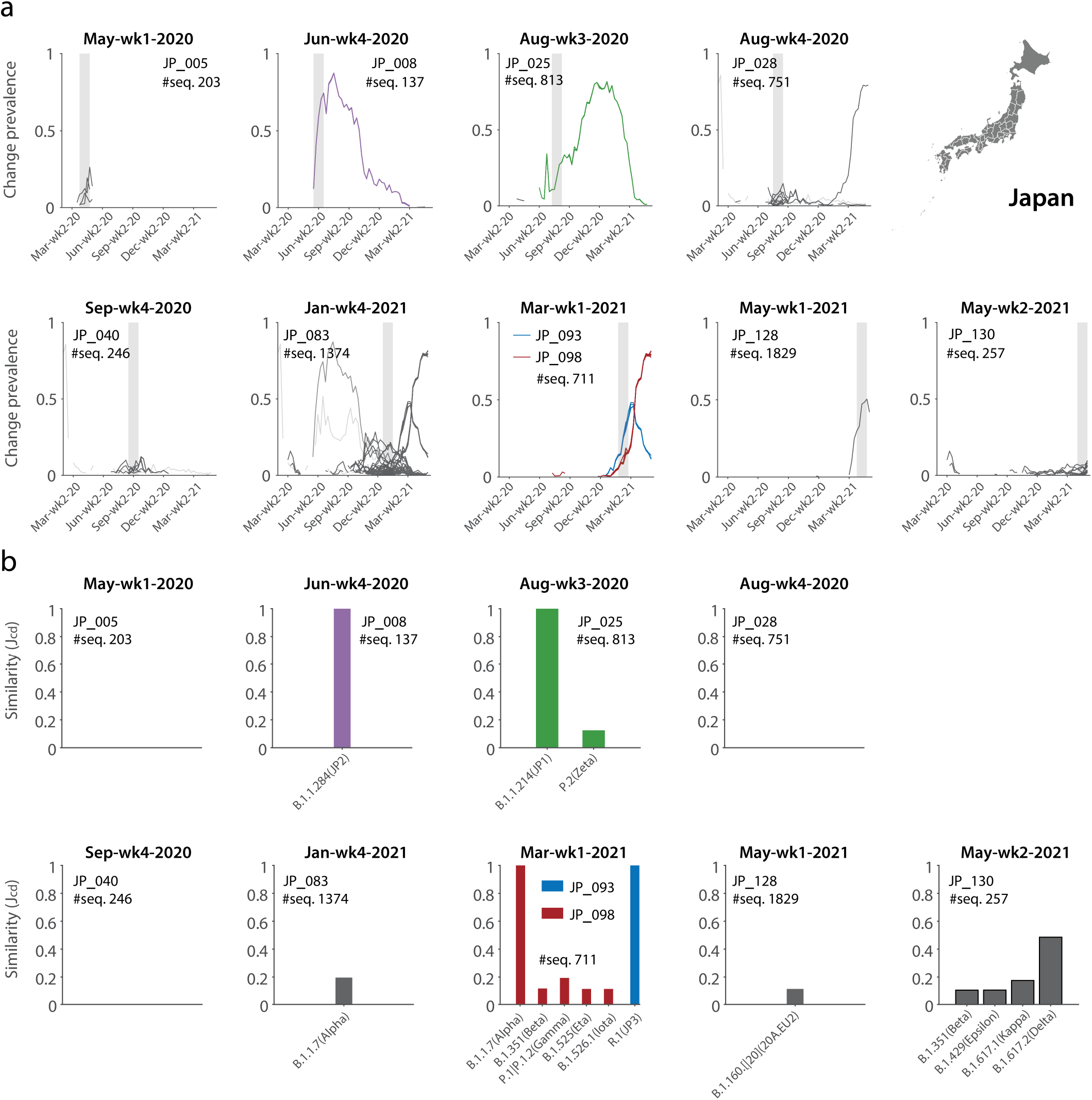
Data-driven warning of possible candidate variant emergence issued in Japan. **a**, The nine time points when warnings were raised. Note that Mar-wk1-21 exhibits two distinct warnings for clusters JP_093 and JP_98. **b**, Cluster-lineage Jaccard similarity for the ten warnings. The four hits (similarity above the threshold *J_cd_* = 0.5) are reported in Fig. 1 in the main text. Strong cluster-lineage similarity is already observed at the time when clustering is performed, which qualifies these as early hits. In this case, candidates were discarded within five weeks, as no late hits (i.e. strong similarity observed with some delay after clustering/warning between a cluster with an amino acid change composition that can be traced back to the original warning and a lineage dictionary) were recorded.

**Extended Data Figure 3.**
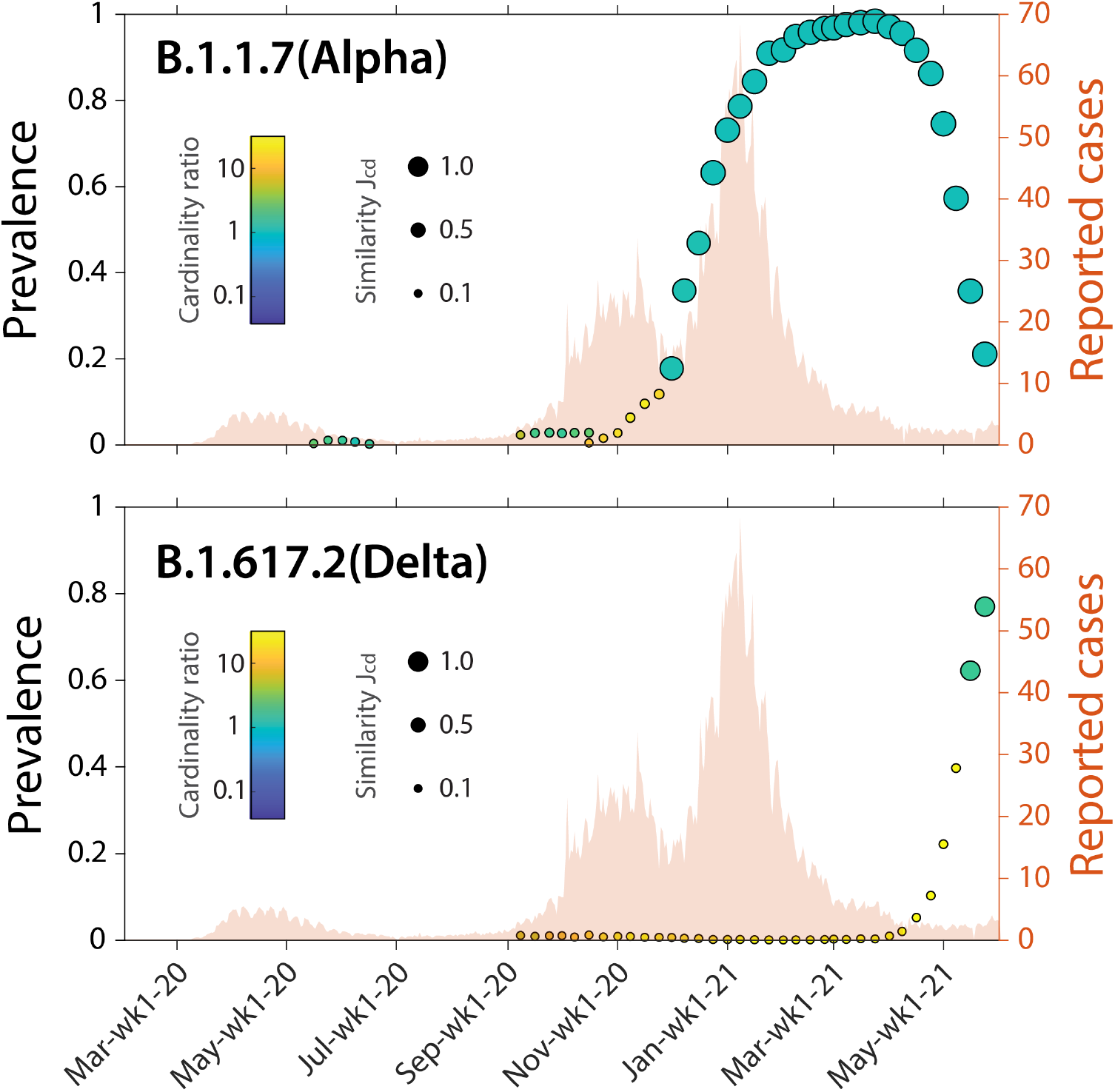
Temporal dynamics of the two variants identified in UK. We show the decline of Alpha variant occurring together with the growth of the Delta variant. Details as in panels d–g of Fig. 1 in the main text.

**Extended Data Figure 4.**
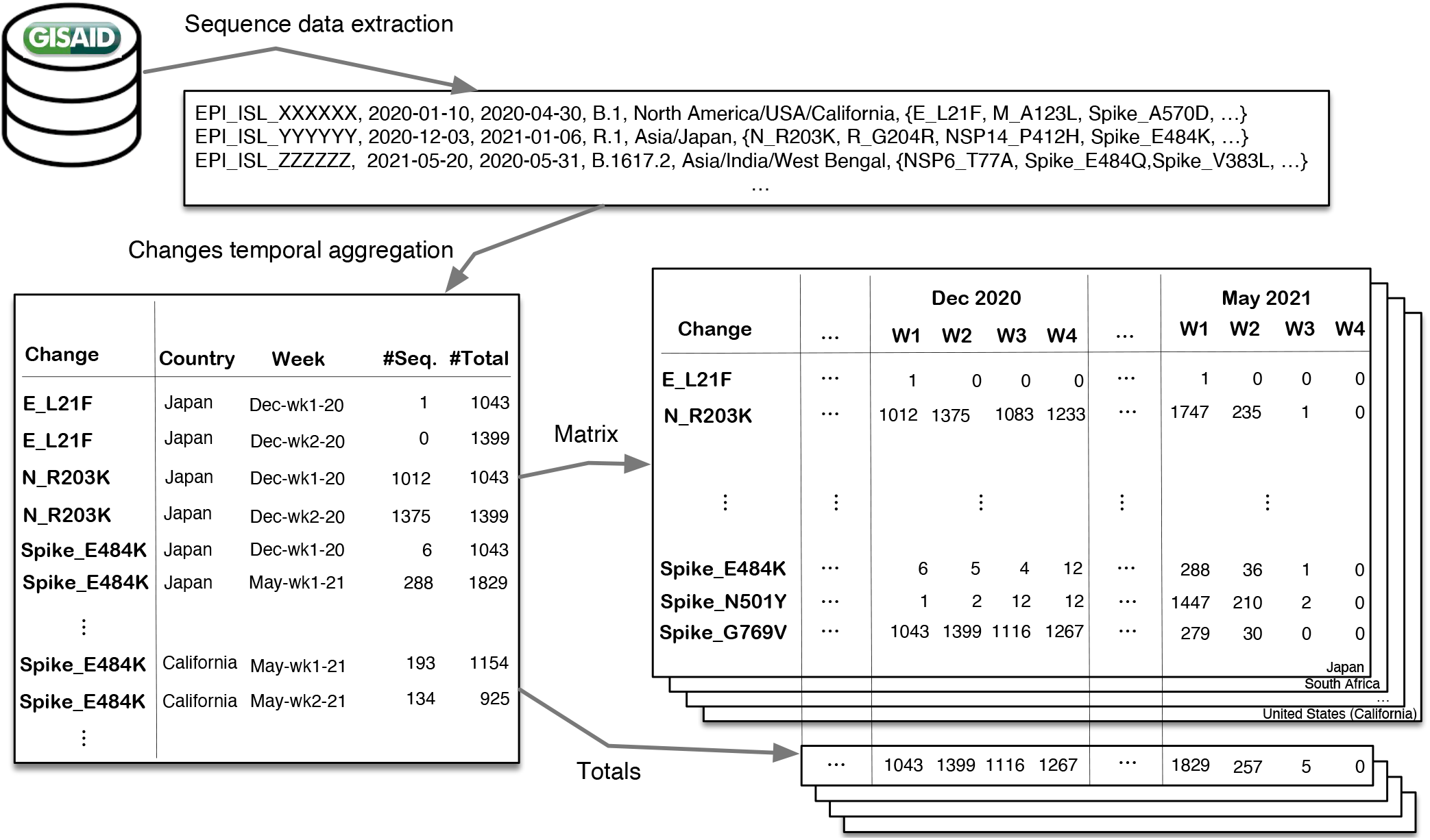
Data extraction and aggregation steps. GISAID original data comprise records of sequence accession ID, collection date, submission date, Pangolin lineage, collection location, and list of amino acid changes. We compute a table of tuples with: amino acid change, country of collection, week of collection, absolute count of sequences holding that change, and total collected sequences in the same country-week. For selected countries (the example shows Japan), we build a matrix where rows are amino acid changes observed in at least five sequences for at least three weeks, columns are observation weeks, and cells contain the number of sequences collected in the given week exhibiting the specific amino acid change. A separate vector holds the total number of sequences collected in the location under study for each of the considered weekly periods. Note that the last weeks of May 2021 hold very few data, thereby producing diluted results of our clustering and warning method.

**Extended Data Table 1.**
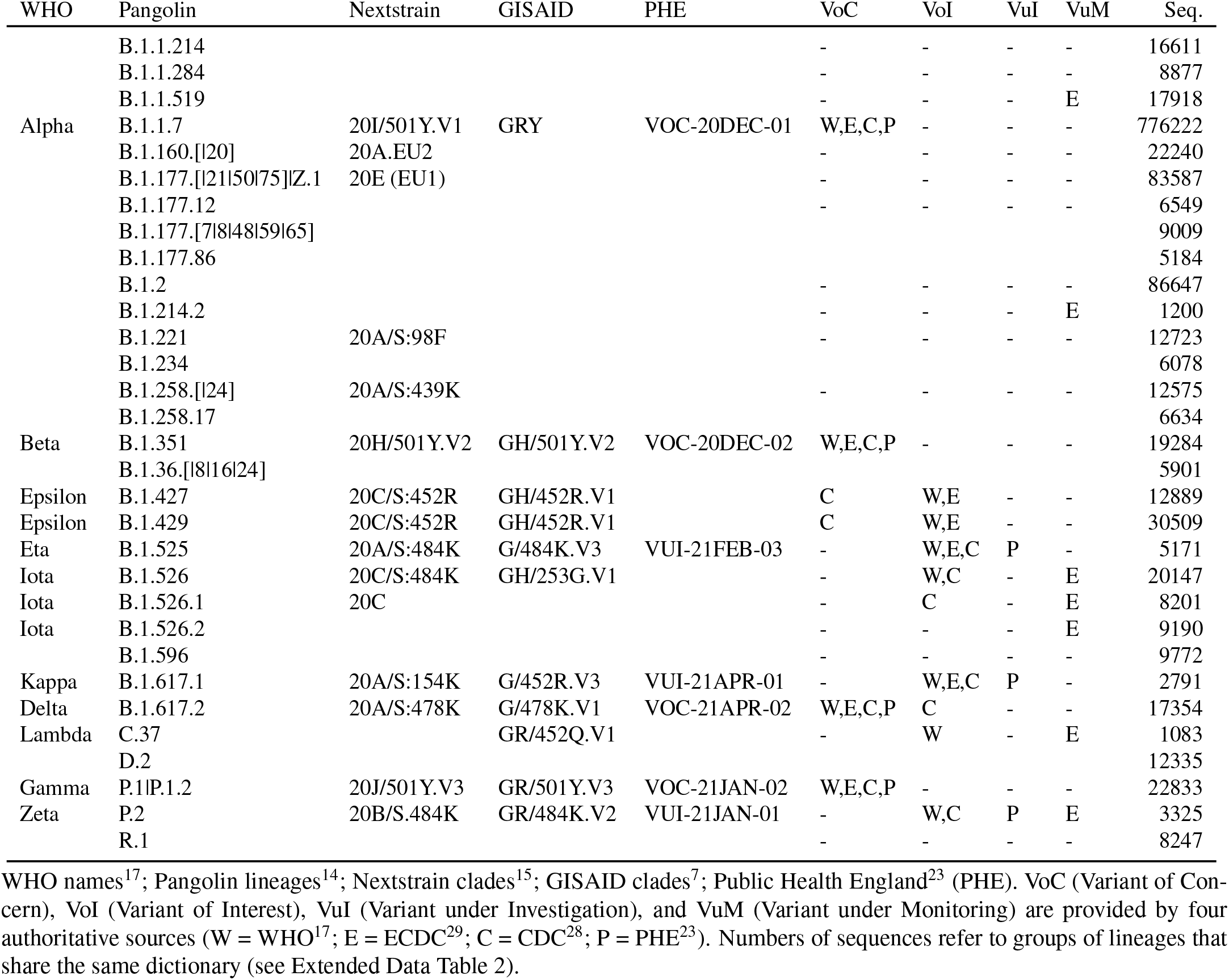
List of considered lineages

**Extended Data Table 2.**
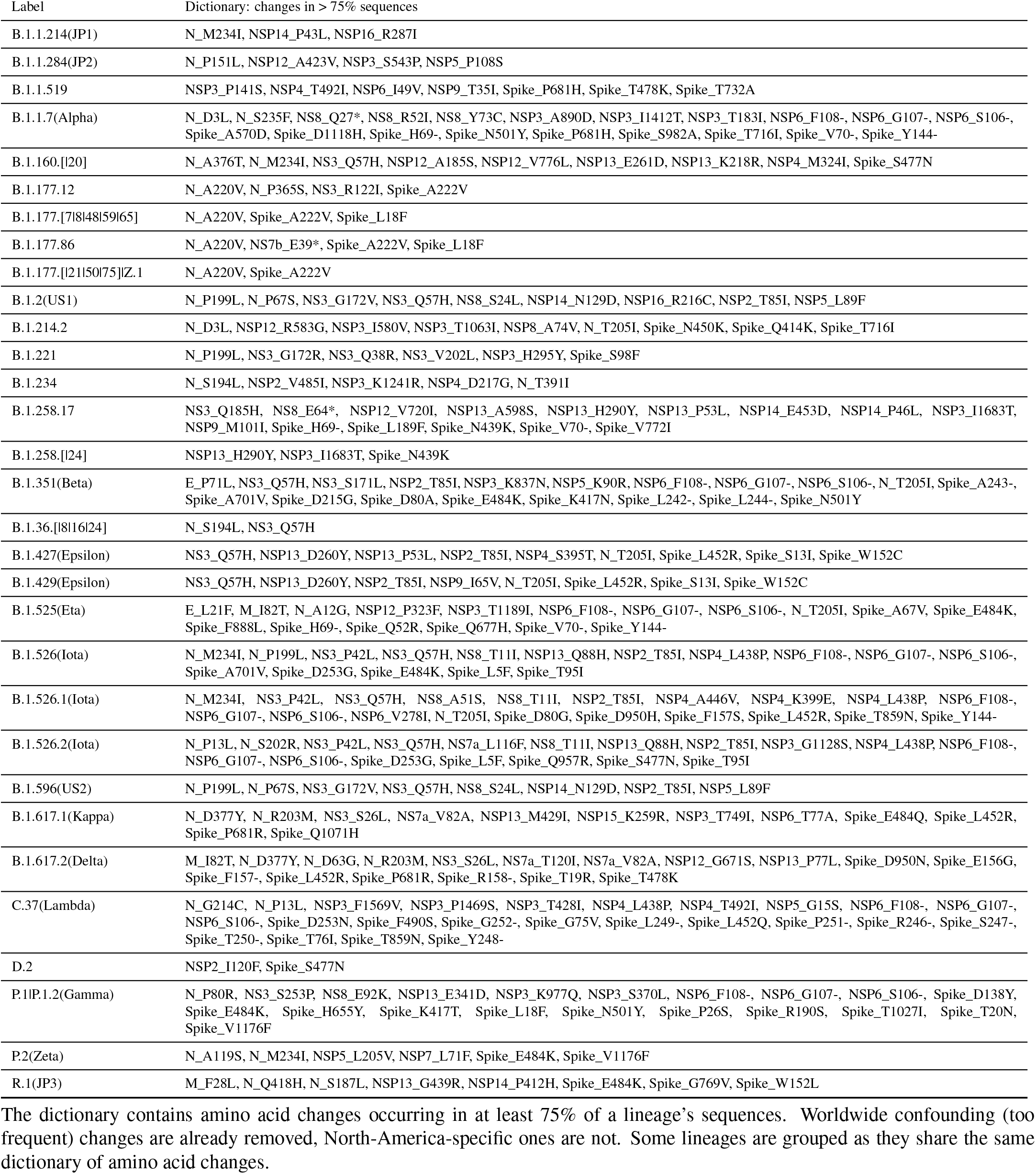
The lineage dictionary

**Extended Data Table 3.** Statistics on SARS-CoV-2 sequencing in world countries

| Country | #Deposited Seq. | COVID-19 Cases | Sequenced% | Delay 2020 | Delay 2021 |
| --- | --- | --- | --- | --- | --- |
| USA | 517,218 | 33,326,436 | 1.55 | 121.29 | 27.70 |
| United Kingdom | 425,455 | 4,515,778 | 9.42 | 37.74 | 15.65 |
| Germany | 119,813 | 3,701,692 | 3.24 | 165.17 | 21.13 |
| Denmark | 102,456 | 284,888 | 35.96 | 55.99 | 29.01 |
| Sweden | 55,158 | 1,076,993 | 5.12 | 88.45 | 46.62 |
| Japan | 49,007 | 755,711 | 6.48 | 130.27 | 50.61 |
| Switzerland | 41,853 | 696,801 | 6.01 | 91.71 | 23.09 |
| France | 39,872 | 6,104,346 | 0.65 | 143.17 | 30.22 |
| Netherlands | 34,314 | 1,684,374 | 2.04 | 61.70 | 24.56 |
| Spain | 30,510 | 3,693,012 | 0.83 | 128.99 | 36.00 |
| Italy | 29,301 | 4,225,163 | 0.69 | 139.11 | 21.65 |
| Canada | 28,000 | 1,395,336 | 2.01 | 178.37 | 73.08 |
| Belgium | 23,278 | 1,066,957 | 2.18 | 76.98 | 18.17 |
| Australia | 17,811 | 30,141 | 59.09 | 67.81 | 12.15 |
| India | 16,309 | 28,574,350 | 0.06 | 111.98 | 46.61 |
| Austria | 15,793 | 645,834 | 2.45 | 116.26 | 50.63 |
| Ireland | 14,316 | 263,252 | 5.44 | 134.10 | 21.79 |
| Poland | 13,245 | 2,874,092 | 0.46 | 109.06 | 24.32 |
| Brazil | 12,863 | 16,803,472 | 0.08 | 162.32 | 46.82 |
| Israel | 11,835 | 839,532 | 1.41 | 48.23 | 62.73 |
| Slovenia | 11,764 | 254,692 | 4.62 | 191.98 | 25.89 |
| Norway | 10,173 | 126,218 | 8.06 | 78.44 | 31.07 |
| Lithuania | 9,914 | 275,545 | 3.60 | 77.22 | 29.60 |
| Mexico | 9,517 | 2,426,822 | 0.39 | 174.72 | 29.03 |
| Portugal | 8,354 | 851,031 | 0.98 | 85.21 | 25.79 |
| Luxembourg | 7,771 | 70,088 | 11.09 | 57.58 | 30.50 |
| South Africa | 7,161 | 1,680,373 | 0.43 | 85.24 | 68.15 |
| Greece | 6,818 | 406,751 | 1.68 | 174.50 | 52.94 |
| South Korea | 6,212 | 142,851 | 4.35 | 105.06 | 33.99 |
| Finland | 5,572 | 92,913 | 6.00 | 154.45 | 63.23 |
| Iceland | 5,070 | 6,555 | 77.35 | 135.17 | 11.46 |
| Turkey | 4,820 | 4,447,074 | 0.11 | 129.20 | 27.69 |
| Philippines | 4,293 | 1,247,899 | 0.34 | 163.09 | 89.50 |
| Russia | 3,968 | 5,040,390 | 0.08 | 108.05 | 57.22 |
| Czech Republic | 3,407 | 1,662,608 | 0.20 | 141.02 | 40.82 |
| Latvia | 3,350 | 134,162 | 2.50 | 119.18 | 34.80 |
| Argentina | 3,218 | 3,884,447 | 0.08 | 196.08 | 44.64 |
| Slovakia | 2,656 | 390,129 | 0.68 | 85.22 | 23.08 |
| Singapore | 2,575 | 62,145 | 4.14 | 89.88 | 10.91 |
| Bulgaria | 2,485 | 419,180 | 0.59 | 133.08 | 40.80 |
| Qatar | 2,256 | 218,080 | 1.03 | 294.87 | 56.50 |
| Chile | 2,199 | 1,403,101 | 0.16 | 164.08 | 25.36 |
| Hong Kong | 1,867 | 11,849 | 15.76 | 125.78 | 64.61 |
| United Arab Emirates | 1,846 | 576,947 | 0.32 | 171.14 | 33.00 |
| Indonesia | 1,845 | 1,837,126 | 0.10 | 129.49 | 63.64 |
| Estonia | 1,782 | 129,909 | 1.37 | 74.25 | 33.12 |
| Croatia | 1,656 | 357,109 | 0.46 | 162.09 | 40.16 |
| Bangladesh | 1,639 | 805,980 | 0.20 | 106.80 | 33.43 |
| Thailand | 1,337 | 169,344 | 0.79 | 99.41 | 25.93 |
| Colombia | 1,334 | 3,488,046 | 0.04 | 144.99 | 40.48 |
| Peru | 1,226 | 1,965,432 | 0.06 | 166.92 | 58.72 |
| China | 1,155 | 90,697 | 1.27 | 117.00 | 16.90 |
| Reunion | 1,094 | - | - | 153.25 | 53.70 |
| New Zealand | 1,064 | 2,682 | 39.67 | 87.06 | 13.84 |

Countries contributing to the GISAID database with more than 1,000 sequences; COVID-19 cases<sup>42</sup>; percentage of sequences over cases; average delay (in days) between sample collection and sequence submission dates.

**Extended Data Table 4.**
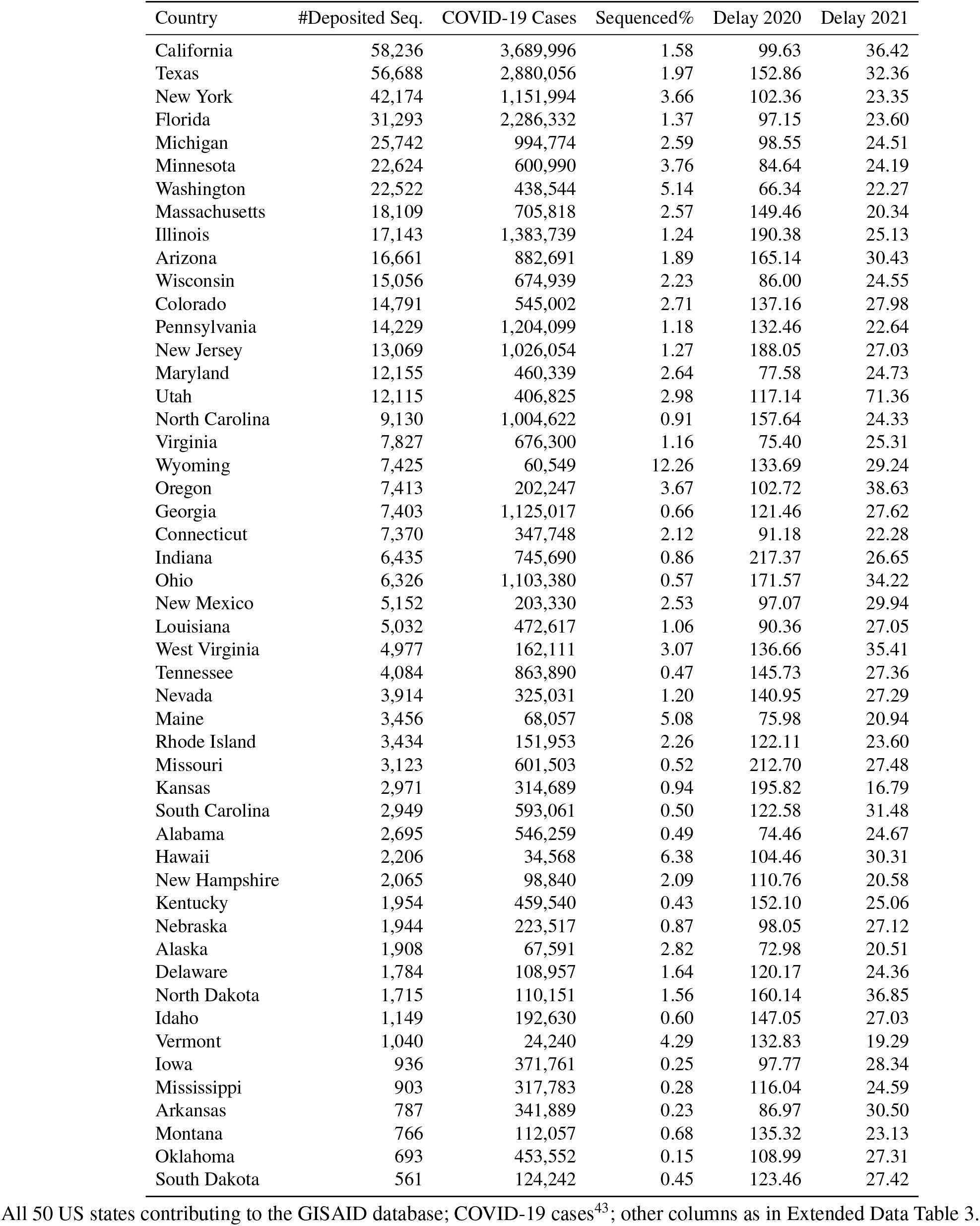
Statistics on SARS-CoV-2 sequencing in all US states

**Extended Data Table 5.**
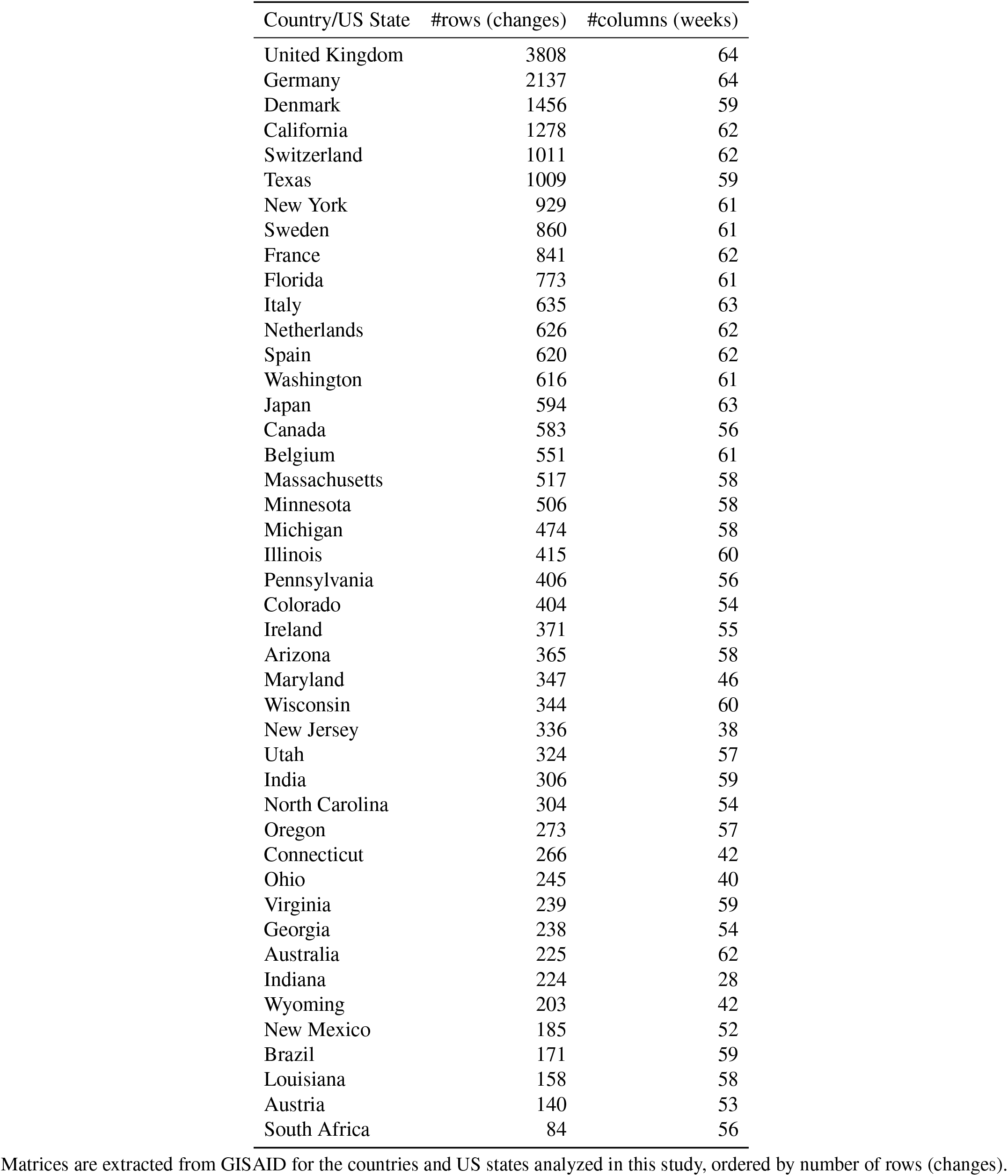
Size of the aggregated data matrices

## References

1. Korber, B. et al. Tracking changes in SARS-CoV-2 Spike: evidence that D614G increases infectivity of the COVID-19 virus. Cell (2020).

2. Wu, A. et al. One year of SARS-CoV-2 evolution. Cell Host & Microbe 29, 503–507 (2021).

3. Sjaarda, C. P. et al. Phylogenomics reveals viral sources, transmission, and potential superinfection in early-stage COVID-19 patients in Ontario, Canada. Scientific Reports 11, 1–9 (2021).

4. Naveca, F. G. et al. COVID-19 in Amazonas, Brazil, was driven by the persistence of endemic lineages and P.1 emergence. Nature Medicine, 1–9 (2021).

5. Alteri, C. et al. Genomic epidemiology of SARS-CoV-2 reveals multiple lineages and early spread of SARS-CoV-2 infections in Lombardy, Italy. Nature Communications 12, 1–13 (2021).

6. Hodcroft, E. B. et al. Spread of a SARS-CoV-2 variant through Europe in the summer of 2020. Nature (2021).

7. Shu, Y. & McCauley, J. GISAID: Global initiative on sharing all influenza data–from vision to reality. Eurosurveillance 22 (2017).

8. Lauring, A. S. & Hodcroft, E. B. Genetic variants of SARS-CoV-2—what do they mean? Jama 325, 529–531 (2021).

9. Ziegler, K., Steininger, P., Ziegler, R., Steinmann, J., Korn, K. & Ensser, A. SARS-CoV-2 samples may escape detection because of a single point mutation in the N gene. Eurosurveillance 25, 2001650 (2020).

10. Wang, R., Hozumi, Y., Yin, C. & Wei, G.-W. Mutations on COVID-19 diagnostic targets. Genomics 112, 5204–5213 (2020).

11. Madhi, S. A. et al. Efficacy of the ChAdOx1 nCoV-19 Covid-19 vaccine against the B.1.351 variant. New England Journal of Medicine 384, 1885–1898 (2021).

12. Planas, D. et al. Sensitivity of infectious SARS-CoV-2 B.1.1.7 and B.1.351 variants to neutralizing antibodies. Nature medicine 27, 917–924 (2021).

13. Garcia-Beltran, W. F. et al. Multiple SARS-CoV-2 variants escape neutralization by vaccine-induced humoral immunity. Cell 184, 2372–2383 (2021).

14. Rambaut, A. et al. A dynamic nomenclature proposal for SARS-CoV-2 lineages to assist genomic epidemiology. Nature Microbiology 5, 1403–1407 (Nov. 2020).

15. Hadfield, J. et al. Nextstrain: real-time tracking of pathogen evolution. Bioinformatics 34, 4121–4123 (Dec. 2018).

16. Callaway, E. Coronavirus variants get Greek names – but will scientists use them? Nature 594, 162 (2021).

17. World Health Organization. Tracking SARS-CoV-2 variants. https://www.who.int/en/activities/tracking-SARS-CoV-2-variants/. (2021). Last accessed: July 3th, 2021.

18. Volz, E. et al. Assessing transmissibility of SARS-CoV-2 lineage B. 1.1. 7 in England. Nature 593, 266–269 (2021).

19. Tegally, H. et al. Detection of a SARS-CoV-2 variant of concern in South Africa. Nature 592, 438–443 (2021).

20. Dhar, M. S. et al. Genomic characterization and Epidemiology of an emerging SARS-CoV-2 variant in Delhi, India. Preprint at https://doi.org/10.1101/2021.06.02.21258076 (2021).

21. Zhang, W., Davis, B. D., Chen, S. S., Sincuir Martinez, J. M., Plummer, J. T. & Vail, E. Emergence of a Novel SARS-CoV-2 Variant in Southern California. JAMA 325, 1324–1326 (2021).

22. Voloch, C. M. et al. Genomic characterization of a novel SARS-CoV-2 lineage from Rio de Janeiro, Brazil. Journal of Virology 95, e00119–21 (2021).

23. Public Health England (PHE). Investigation of SARS-CoV-2 variants of concern: technical briefings. https://www.gov.uk/government/publications/investigation-of-novel-sars-cov-2-variant-variant-of-concern-20201201. (2021). Last accessed: July 3th, 2021.

24. Tablizo, F. A. et al. Genome sequencing and analysis of an emergent SARS-CoV-2 variant characterized by multiple spike protein mutations detected from the Central Visayas Region of the Philippines. Preprint at https://doi.org/10.1101/2021.03.03.21252812 (2021).

25. West Jr, A. P., Barnes, C. O., Yang, Z. & Bjorkman, P. J. SARS-CoV-2 lineage B.1.526 emerging in the New York region detected by software utility created to query the spike mutational landscape. Preprint at https://doi.org/10.1101/2021.02.14.431043 (2021).

26. Cherian, S. et al. Convergent evolution of SARS-CoV-2 spike mutations, L452R, E484Q and P681R, in the second wave of COVID-19 in Maharashtra, India. Preprint at https://doi.org/10.1101/2021.04.22.440932 (2021).

27. Romero, P. E. et al. Novel sublineage within B.1.1.1 currently expanding in Peru and Chile, with a convergent deletion in the ORF1a gene (Δ3675-3677) and a novel deletion in the Spike gene (Δ246-252, G75V, T76I, L452Q, F490S, T859N). https://virological.org/t/novel-sublineage-within-b-1-1-1-currently-expanding-in-peru-and-chile-with-a-convergent-deletion-in-the-orf1a-gene-3675-3677-and-a-novel-deletion-in-the-spike-gene-246-252-g75v-t76i-l452q-f490s-t859n/685. (2021). Last accessed: July 3th, 2021.

28. Centers for Disease Control and Prevention. SARS-CoV-2 Variant Classifications and Definitions. https://www.cdc.gov/coronavirus/2019-ncov/variants/variant-info.html. (2021). Last accessed: July 3th, 2021.

29. European Centre for Disease Prevention and Control. SARS-CoV-2 variants of concern. https://www.ecdc.europa.eu/en/covid-19/variants-concern. (2021). Last accessed: July 3th, 2021.

30. Bernasconi, A. et al. VirusViz: comparative analysis and effective visualization of viral nucleotide and amino acid variants. Nucleic Acids Research (2021).

31. Mercatelli, D. & Giorgi, F. M. Geographic and genomic distribution of SARS-CoV-2 mutations. Frontiers in Microbiology 11, 1800 (2020).

32. Wang, R., Chen, J., Gao, K., Hozumi, Y., Yin, C. & Wei, G.-W. Analysis of SARS-CoV-2 mutations in the United States suggests presence of four substrains and novel variants. Communications Biology 4, 1–14 (2021).

33. Troyano-Hernáez, P., Reinosa, R. & Holguín, Á. Evolution of SARS-CoV-2 envelope, membrane, nucleocapsid, and spike structural proteins from the beginning of the pandemic to September 2020: a global and regional approach by epidemiological week. Viruses 13, 243 (2021).

34. Chiara, M., Horner, D. S., Gissi, C. & Pesole, G. Comparative Genomics Reveals Early Emergence and Biased Spatiotemporal Distribution of SARS-CoV-2. Molecular Biology and Evolution 38, 2547–2565 (2021).

35. Yang, H.-C. et al. Analysis of genomic distributions of SARS-CoV-2 reveals a dominant strain type with strong allelic associations. Proceedings of the National Academy of Sciences 117, 30679–30686 (2020).

36. Wada, K., Wada, Y. & Ikemura, T. Time-series analyses of directional sequence changes in SARS-CoV-2 genomes and an efficient search method for candidates for advantageous mutations for growth in human cells. Gene: X 5, 100038 (2020).

37. Showers, W. M., Leach, S. M., Kechris, K. & Strong, M. Analysis of SARS-CoV-2 Mutations Over Time Reveals Increasing Prevalence of Variants in the Spike Protein and RNA-Dependent RNA Poly-merase. Preprint at https://doi.org/10.1101/2021.03.05.433666 (2021).

38. O’Toole, Á. et al. Tracking the international spread of SARS-CoV-2 lineages B.1.1.7 and B.1.351/501Y-V2. Wellcome Open Research 6, 121 (2021).

39. Wall, E. C. et al. Neutralising antibody activity against SARS-CoV-2 VOCs B.1.617.2 and B.1.351 by BNT162b2 vaccination. The Lancet 397, 2331–2333 (2021).

40. Grubaugh, N. D., Hodcroft, E. B., Fauver, J. R., Phelan, A. L. & Cevik, M. Public health actions to control new SARS-CoV-2 variants. Cell 184, 1127–1132 (2021).

41. Fruchterman, T. M. & Reingold, E. M. Graph drawing by force-directed placement. Software: Practice and Experience 21, 1129–1164 (1991).

42. Ritchie, H. et al. Coronavirus Pandemic (COVID-19). https://ourworldindata.org/coronavirus. (2020). Last accessed: July 3th, 2021.

43. Centers for Disease Control and Prevention. United States COVID-19 Cases and Deaths by State over Time. https://catalog.data.gov/dataset/united-states-covid-19-cases-and-deaths-by-state-over-time. (2021). Last accessed: July 3th, 2021.

## References

44. Canakoglu, A., Pinoli, P., Bernasconi, A., Alfonsi, T., Melidis, D. P. & Ceri, S. ViruSurf: an integrated database to investigate viral sequences. Nucleic Acids Research 49, D817–D824 (2020).

45. Mullen, J. L. et al. Outbreak.info. https://outbreak.info/. (2020). Last accessed: July 3th, 2021.

46. Hodcroft, E. B. CoVariants: SARS-CoV-2 Mutations and Variants of Interest. https://covariants.org/. (2021). Last accessed: July 3th, 2021.

47. Zhou, P. et al. A pneumonia outbreak associated with a new coronavirus of probable bat origin. Nature 579, 270–273 (2020).

48. Kaufman, L. & Rousseeuw, P. J. Finding Groups in Data: an Introduction to Cluster Analysis (John Wiley & Sons, 2009).

49. Sakoe, H. & Chiba, S. Dynamic programming algorithm optimization for spoken word recognition. IEEE Transactions on Acoustics, Speech, and Signal Processing 26, 43–49 (1978).

50. Rousseeuw, P. J. Silhouettes: a graphical aid to the interpretation and validation of cluster analysis. Journal of Computational and Applied Mathematics 20, 53–65 (1987).

51. Kendall, M. G. Rank Correlation Methods (Griffin, 1948).

52. Burkey, J. Mann-Kendall Tau-b with Sen’s Method (enhanced). MATLAB Central File Exchange. https://www.mathworks.com/matlabcentral/fileexchange/11190-mann-kendall-tau-b-with-sen-s-method-enhanced. (2021). Retrieved: April 1st, 2021.

53. Jaccard, P. Étude comparative de la distribution florale dans une portion des Alpes et des Jura. Bull Soc Vaudoise Sci Nat 37, 547–579 (1901).

54. Rambaut, A. et al. Preliminary genomic characterisation of an emergent SARS-CoV-2 lineage in the UK defined by a novel set of spike mutations. https://virological.org/t/preliminary-genomic-characterisation-of-an-emergent-sars-cov-2-lineage-in-the-uk-defined-by-a-novel-set-of-spike-mutations/563. (2020). Last accessed: July 3th, 2021.

55. Health Department - Republic of South Africa. COVID-19 South African Online Portal. Update on Covid-19 (18th December 2020). https://sacoronavirus.co.za/2020/12/18/update-on-covid-19-18th-december-2020/. (2020). Last accessed: July 3th, 2021.

56. Tegally, H. et al. Emergence and rapid spread of a new severe acute respiratory syndrome-related coronavirus 2 (SARS-CoV-2) lineage with multiple spike mutations in South Africa. Preprint at https://doi.org/10.1101/2020.12.21.20248640 (2020).

57. National Institute of Infection Diseases (NIID) of Japan. Brief report: New Variant Strain of SARS-CoV-2 Identified in Travelers from Brazil. https://www.niid.go.jp/niid/en/2019-ncov-e/10108-covid19-33-en.html. (2021). Last accessed: July 3th, 2021.

58. Naveca, F. et al. Phylogenetic relationship of SARS-CoV-2 sequences from Amazonas with emerging Brazilian variants harboring mutations E484K and N501Y in the Spike protein. https://virological.org/t/phylogenetic-relationship-of-sars-cov-2-sequences-from-amazonas-with-emerging-brazilian-variants-harboring-mutations-e484k-and-n501y-in-the-spike-protein/585. (2021). Last accessed: July 3th, 2021.

59. Faria, N. R. et al. Genomic characterisation of an emergent SARS-CoV-2 lineage in Manaus: preliminary findings. https://virological.org/t/genomic-characterisation-of-an-emergent-sars-cov-2-lineage-in-manaus-preliminary-findings/586. (2021). Last accessed: July 3th, 2021.

60. California Department of Public Health. COVID-19 Variant First Found in Other Countries and States Now Seen More Frequently in California. https://www.cdph.ca.gov/Programs/OPA/Pages/NR21-020.aspx. (2021). Last accessed: July 3th, 2021.

61. Zhang, W., Davis, B. D., Chen, S. S., Sincuir Martinez, J. M., Plummer, J. T. & Vail, E. Emergence of a novel SARS-CoV-2 strain in Southern California, USA. Preprint at https://doi.org/10.1101/2021.01.18.21249786 (2021).

62. Voloch, C. M. et al. Genomic characterization of a novel SARS-CoV-2 lineage from Rio de Janeiro, Brazil. Preprint at https://doi.org/10.1101/2020.12.23.20248598 (2020).

63. Annavajhala, M. K. et al. A novel SARS-CoV-2 variant of concern, B.1.526, identified in New York. Preprint at https://doi.org/10.1101/2021.02.23.21252259 (2021).

64. Ministry of Health and Family Welfare, India. Genome Sequencing by INSACOG shows variants of concern and a Novel variant in India. https://pib.gov.in/PressReleaseIframePage.aspx?PRID=1707177. (2021). Last accessed: July 3th, 2021.

65. Bernini, A., Toure, A. L. & Casagrandi, R. The time varying network of urban space uses in Milan. Applied Network Science 2019, 128 (2019).

66. Blondel, V. D., Guillaume, J.-L., Lambiotte, R. & Lefebvre, E. Fast unfolding of communities in large networks. Journal of Statistical Mechanics: Theory and Experiment 2008, P10008 (2008).

67. Kehagias, A. Community Detection Toolbox. MATLAB Central File Exchange. https://www.mathworks.com/matlabcentral/fileexchange/45867-community-detection-toolbox. (2021). Retrieved: April 1st, 2021.

